# Multiple laboratory mouse reference genomes define strain specific haplotypes and novel functional loci

**DOI:** 10.1101/235838

**Authors:** Jingtao Lilue, Anthony G. Doran, Ian T. Fiddes, Monica Abrudan, Joel Armstrong, Ruth Bennett, William Chow, Joanna Collins, Stephan Collins, Anne Czechanski, Petr Danecek, Mark Diekhans, Dirk-Dominic Dolle, Matt Dunn, Richard Durbin, Dent Earl, Anne Ferguson-Smith, Paul Flicek, Jonathan Flint, Adam Frankish, Beiyuan Fu, Mark Gerstein, James Gilbert, Leo Goodstadt, Jennifer Harrow, Kerstin Howe, Mikhail Kolmogorov, Stefanie Köenig, Chris Lelliott, Jane Loveland, Clayton E. Mathews, Richard Mott, Paul Muir, Fabio Navarro, Duncan Odom, Naomi Park, Sarah Pelan, Son K Phan, Michael Quail, Laura Reinholdt, Lars Romoth, Lesley Shirley, Cristina Sisu, Marcela Sjoberg-Herrera, Mario Stanke, Charles Steward, Mark Thomas, Glen Threadgold, David Thybert, James Torrance, Kim Wong, Jonathan Wood, Binnaz Yalcin, Fengtang Yang, David J. Adams, Benedict Paten, Thomas M. Keane

## Abstract

The most commonly employed mammalian model organism is the laboratory mouse. A wide variety of genetically diverse inbred mouse strains, representing distinct physiological states, disease susceptibilities, and biological mechanisms have been developed over the last century. We report full length draft *de novo* genome assemblies for 16 of the most widely used inbred strains and reveal for the first time extensive strain-specific haplotype variation. We identify and characterise 2,567 regions on the current Genome Reference Consortium mouse reference genome exhibiting the greatest sequence diversity between strains. These regions are enriched for genes involved in defence and immunity, and exhibit enrichment of transposable elements and signatures of recent retrotransposition events. Combinations of alleles and genes unique to an individual strain are commonly observed at these loci, reflecting distinct strain phenotypes. Several immune related loci, some in previously identified QTLs for disease response have novel haplotypes not present in the reference that may explain the phenotype. We used these genomes to improve the mouse reference genome resulting in the completion of 10 new gene structures, and 62 new coding loci were added to the reference genome annotation. Notably this high quality collection of genomes revealed a previously unannotated gene *(Efcab3-like)* encoding 5,874 amino acids, one of the largest known in the rodent lineage. Interestingly, *Efcab3-like^−/−^* mice exhibit severe size anomalies in four regions of the brain suggesting a mechanism of *Efcab3-like* regulating brain development.

## Background

For over a century inbred laboratory mice have excelled as the premier organism to investigate the genetic basis of morphological, physiological, behavioural and genetic variation^1–3^. The inbred laboratory mouse, C57BL/6J, was the second mammalian genome after human to be fully sequenced, underscoring the prominence of the mouse as a model for mammalian biology^4,5^. The generation and assembly of a reference genome for C57BL/6J accelerated the discovery of the genetic landscape underlying phenotypic variation^4,5^. Inbred laboratory strains are broadly organised into two groups; classical and wild-derived strains, a phenotypically and genetically diverse cohort capturing high allelic diversity, that can be used to model the variation observed in human populations^6,7^. Inbred laboratory strains of wild-derived origin represent a rich source of differential phenotypic responses and genetic diversity not present in classical strains. Wild-derived strains contain between 4 and 8 times more SNPs relative to the reference genome than classical strains^8^ and comprise of genetically distinct subspecies of *Mus musculus* (e.g. *Mus musculus castaneus, Mus musculus musculus)* and *Mus spretus,* that unequivocally contain many unique alleles including novel resistance and susceptibility haplotypes of relevance to human health^9–11^. Several multiparent populations have been derived from inbred laboratory strains to create powerful resources in which to perform genetic mapping of phenotypes and traits^12^. In particular the Collaborative Cross (CC) and Diversity Outbred Cross (DO) have harnessed the extreme genetic and phenotypic diversity of the wild-derived strains to map a range of phenotypes including benzene toxicity^13^ and immune responses to influenza^14^, Ebola^15^ and allergic airway inflammation^16^.

Next-generation sequencing enabled the creation of genome-wide variation catalogues (SNPs, short indels, and structural variation) for thirty-six laboratory mouse strains and facilitated the identification of strain specific mutations^8,17^, interpretation of variants’ effects on gene expression and gene regulation^8^, the genetic architecture of complex traits^18^, and the detection of regions of the greatest genetic divergence from the reference strain^19,20^. However, reliance on mapping next-generation sequencing reads to a single reference genome (C57BL/6J) has meant that the true extent of strain specific variation is unknown. In some strain comparisons, the difference between reference and sequenced strain genomes is as large as that between human and chimpanzee, making it hard to distinguish whether a read is mis-mapped or highly divergent. *De novo* genome assembly methods overcome the limitations of mapping back to a reference genome, thereby allowing unbiased assessments of the differences between genomes, thus providing improved catalogues of all classes of variants. Not only are these resources essential for understanding the relationship between genetic and phenotypic variation, but the ability to investigate sequence variation across different evolutionary time scales provides a powerful tool to explore functional annotations^21^.

We have completed the first draft *de novo* assembled genome sequences and strain specific gene annotation of twelve classical inbred laboratory strains (129S1/SvImJ, A/J, AKR/J, BALB/cJ, C3H/HeJ, C57BL/6NJ, CBA/J, DBA/2J, FVB/NJ, LP/J, NZO/HlLtJ and NOD/ShiLtJ) and four wild-derived strains representing the *Mus; M. m. castaneus* (CAST/EiJ), *M. m. musculus* (PWK/PhJ), *M. m. domesticus* (WSB/EiJ) and *M. spretus* (SPRET/EiJ) backgrounds. This collection comprises a large and diverse array of laboratory strains, including those closely related to commonly used mouse cell lines (BALB/3T3 and L929, derived from BALB/c and C3H related strains), embryonic stem cell derived gene knockouts (historically 129-related strains)^22^, humanised mouse models (primarily NOD-related nude mice)^1^, gene knockout background strains (C57BL/6NJ)^23^, the founders of commonly used recombinant inbred lines such as the AKXD, BXA, BXD, CXB and CC^24^, and outbred mapping populations such as the DO and the heterogeneous stock (HS)^25^.

We combined previous variation catalogs^8,26^ and the assembled genome sequences to identify regions of greatest haplotype diversity, and found that these regions are enriched for genes associated with immunity, sensory perception, behaviour and kin recognition. These regions are statistically enriched for young transposable elements, adding to increasing evidence that transposons play a significant role in generating haplotype diversity in mice. The strain assemblies revealed many novel gene family member combinations, novel alleles not found in the reference genome, and highlight previously unknown levels of inter-strain variation. Complementary techniques were used to confirm novel gene family content and composition revealed by the assemblies, including loci notable for potential roles in hybrid sterility, environmental sensing and pathogen response. We improved both sequence and gene annotation of the C57BL/6J mouse reference, and identify novel genes including a previously unannotated 188 exon (5,874 amino acid) gene present in all strains and is conserved across many mammalian species, and plays a role in the development of several mouse brain regions.

## Results

### Sequence assemblies and genome annotation

Chromosome scale assemblies were produced for 16 laboratory mouse strains using a mixture of Illumina paired-end (40-70x), mate-pairs (3, 6, 10Kbp), fosmid and BAC-end sequences (Supplementary Table 1). For PWK/PhJ, CAST/EiJ, and SPRET/EiJ, Dovetail Genomics Chicago^™^ libraries^27^ were used to provide additional long-range accuracy and sequence contiguity. Pseudo-chromosomes were produced in parallel utilising cross-species synteny alignments to guide assembly resulting in genome assemblies between 2.254 (WSB/EiJ) to 2.328 Gbps (AKR/J) excluding unknown scaffold gap bases. Approximately 0.5-2% of the total genome length per strain was not placed onto a chromosome. The unplaced sequence contigs are composed primarily of unknown gap bases (18-49%) and repeat sequences (61-79%) (Supplementary Table 2), and contain between 89-410 predicted genes per strain (Supplementary Table 3). Mitochondrial genome (mtDNA) assemblies for 14 strains support previously published sequences^28^, although a small number of high quality novel sequence variants in AKR/J, BALB/cJ, C3H/HeJ, and LP/J conflicted with GenBank entries (Supplementary Table 4). Novel mtDNA haplotypes were identified in PWK/PhJ and NZO/HlLtJ. Notably, NZO/HlLtJ contains 55 SNPs (33 shared with the wild-derived strains) and appears distinct compared to the other classical inbred strains (Supplementary Figure 1). Previous variation catalogues have indicated high concordance (>97%) between NZO/HlLtJ and another inbred laboratory strain NZB/BlNJ^19^.

We assessed the base accuracy of the strain chromosomes relative to two versions of the C57BL/6J reference genome (MGSCv3^4^ and GRCm38^5^) by first realigning all of the paired-end sequencing reads from each strain back to the respective assembled genome and identifying SNPs and indels. The combined SNP and indel error rate was between 0.09-0.1 errors per Kbp, compared to 0.334 for MGSCv3 and 0.02 for GRCm38 (Supplementary Table 5). Next we used a set of 612 PCR primer pairs previously used to validate structural variant calls in eight of the laboratory strains^29^. The assemblies had between 4.7-6.7% primer pairs with incorrect alignment compared to 10% for MGSCv3 (Supplementary Table 6). Finally, using PacBio long read cDNA sequencing from liver and spleen of C57BL/6J, CAST/EiJ, PWK/PhJ, and SPRET/EiJ, concordant alignment rates were determined. In both tissues, the GRCm38 reference genome had the highest proportion of correctly aligned cDNA reads (63.1 and 69.8%, respectively) (Supplementary Table 7). The representation of known mouse repeat families in the assemblies shows that short repeat (<200bp) content is comparable to GRCm38 (Supplementary Figure 2a), including short repeats diverged from their respective transposable element (TE) consensus sequence (Supplementary Figure 2b). The total number of long repeat types (>200bp) is consistent across all strains, however the total sequence lengths are consistently shorter than GRCm38 indicating underrepresentation of long repeat sequences (Supplementary Figure 2c).

Strain specific consensus gene sets were produced using two sources of evidence, the GENCODE C57BL/6J annotation, and strain specific RNA-Seq from multiple tissues^30^ (Supplementary Table 8, Supplementary Figure 3). Per strain, the consensus gene sets contain over 20,000 protein coding genes and over 18,000 non-coding genes (Figure 1a, Supplementary Table 1). For the classical laboratory strains 90.2% of coding transcripts (88.0% in wild-derived strains) and 91.2% of noncoding transcripts (91.4% in wild-derived strains) present in the GRCm38 reference gene set were comparatively annotated. Gene predictions from strain specific RNA-Seq (Comparative Augustus^31^) added an average of 1,400 new isoforms to wild-derived and 1,207 new isoforms to classical strain gene annotation sets. Gene prediction based on PacBio cDNA sequencing introduced an average of 1,865 further new isoforms to CAST/EiJ, PWK/PhJ and SPRET/EiJ. Putative novel loci are defined as spliced genes that were predicted from strain specific RNA-Seq and did not overlap any genes projected from the reference genome. On average, 37 genes are putative novel loci (Supplementary Data 1) in wild-derived strains, and 22 in classical strains. Most often these appear to result from gene duplication events. Additionally, an automated pseudogene annotation workflow, Pseudopipe^32^, alongside manually curated pseudogenes lifted-over from the GRCm38 reference genome, identified an average of approximately 11,000 (3,317 conserved between all strains) pseudogenes per strain (Supplementary Figure 4) that appear to have arisen either through retrotransposition (∼l80%) or gene duplication events (∼ 20%).

**Figure 1:**
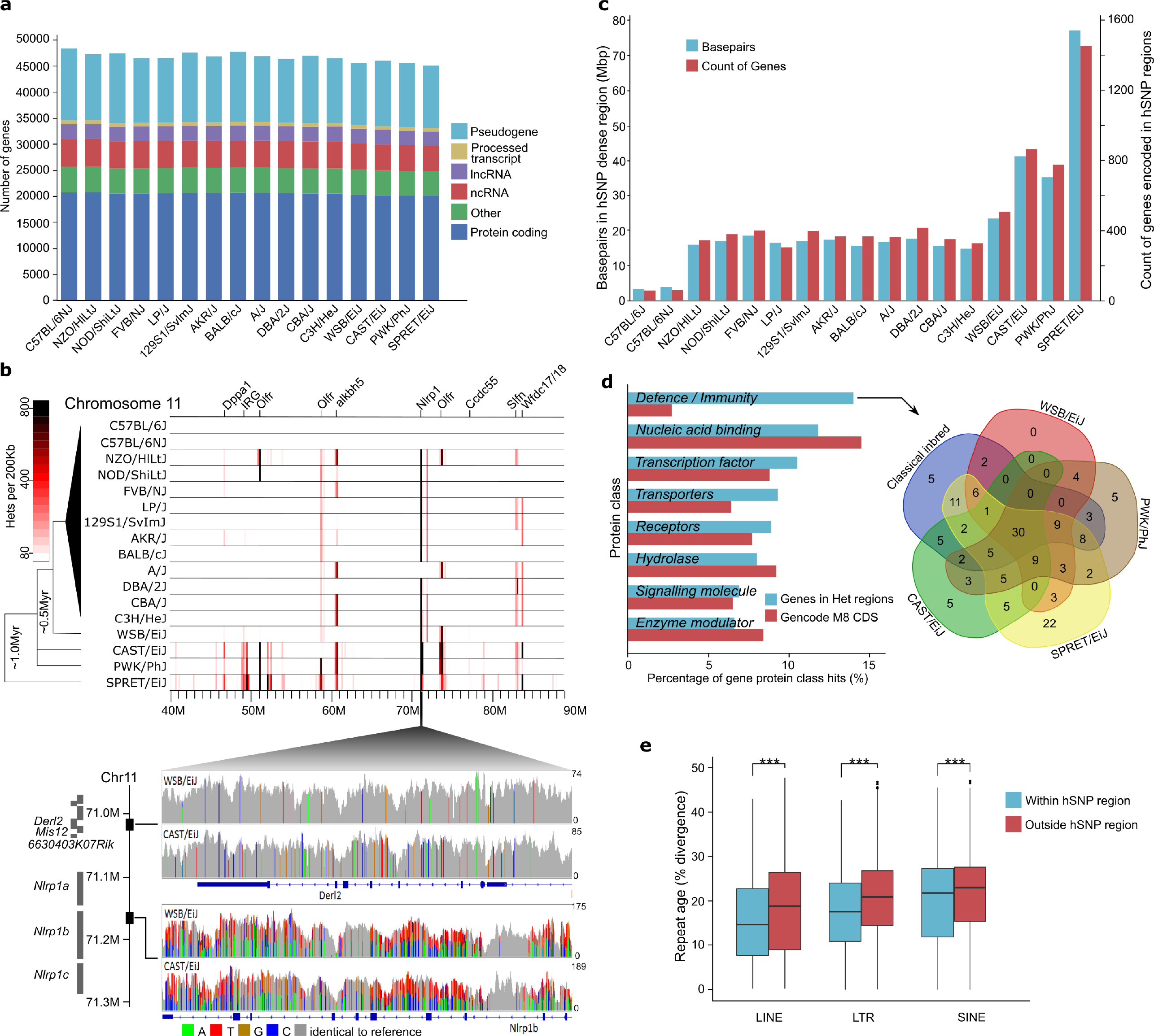
**(a)** Summary of the strain specific gene sets showing the number of genes broken down by GENCODE biotype. **(b)** Heterozygous SNP density for a 50Mbp interval on chromosome 11 in 200Kbp windows for 17 inbred mouse strains based on sequencing read alignments to the C57BL/6J (GRCm38) reference genome (top). Labels indicate genes overlapping the most dense regions. SNPs visualized in CAST/EiJ and WSB/EiJ for 71.006-71.170Mbp on GRCm38 (bottom), including *Derl2,* and *Mis12* (upper panel) and *Nlrp1b* (lower panel). Grey indicates the strain base agrees with the reference, other colours indicate SNP differences, and height corresponds to sequencing depth. **(c)** Total amount of sequence and protein coding genes in regions enriched for heterozygous SNPs per strain. **(d)** Top PantherDB categories of coding genes in regions enriched for heterozygous SNPs based on protein class (left). Intersection of genes in the defence/immunity category for the wild-derived and classical inbred strains (right). **(e)** Box plot of sequence divergence (%), for LTRs, LINEs and SINEs within and outside of heterozygous dense regions. Sequence divergence is relative to a consensus sequence for the transposable element type.

### Regions of the mouse genome exhibiting extreme allelic variation

Inbred laboratory mouse strains are characterized by at least twenty generations of inbreeding, and are genetically homozygous at almost all loci^2,28^. Despite this, previous SNP variation catalogs have identified high quality heterozygous SNPs (hSNPs) when reads are aligned to the C57BL/6J reference genome^33^, and are often removed as genotyping errors. The presence of higher densities of hSNPs may indicate copy number changes, or novel genes that are not present in the reference assembly, forced to partially map to a single locus in the reference^8,19^. Thus their identification and interpretation is a powerful tool for finding errors in a genome assembly and novel, previously unrecognized features. We identified between 116,439 (C57BL/6NJ) and 1,895,741 (SPRET/EiJ) high quality hSNPs from the MGP variation catalogue v5^19^ (Supplementary Table 9). We focused our analysis on the top 5% most hSNP dense regions (windows >= 71 hSNPs per 10Kbp sliding window). This successfully identified the majority of known polymorphic regions among the strains (Supplementary Figures 5) and accounted for ∼49% of all hSNPs available (Supplementary Table 9, Supplementary Figure 6a). After applying this cut-off to all strain-specific hSNP regions and merging overlapping or adjacent windows, between 117 (C57BL/6NJ) and 2,567 (SPRET/EiJ) hSNP regions remained per strain (Supplementary Figure 6b). Many hSNP clusters overlap immunity (e.g. MHC, NOD-like receptors and AIM-like receptors), sensory (e.g. olfactory and taste receptors), reproductive (e.g pregnancy specific glycoproteins and Sperm-associated E-Rich proteins), neuronal and behavior related genes (e.g. itch receptors^34^ and γ-protocadherins^35^) (Figure 1b, Supplementary Figure 5). The wild-derived strain, SPRET/EiJ contained the largest number of genes (1,442) and protein coding sequence base pairs (1.2Mbp) within hSNP dense regions (Figure 1c). All of the wild-derived strains contained gene and CDS base pair counts larger than any classical inbred strain (≥503 and ≥0.36Mbp, respectively; Supplementary Table 9). Notably, the regions identified in C57BL/6J and C57BL/6NJ (117 and 141, respectively; 145 combined) intersect known GRCm38 assembly issues including gaps, unplaced scaffolds or centromeric regions (104/145, 71.7%). The remaining candidate regions include large protein families (15/145, 10.3%) and repeat elements (17/145, 11.7%) (Supplementary Data 2).

We examined protein classes present in the hSNP regions by identifying 1,109 PantherDB matches, assigned to 26 protein classes, from a combined set of all genes in hSNP dense regions (Supplementary Data 3). Defence and immunity was the largest represented protein class in our gene set (155 genes, Supplementary Data 4), accounting for (13.98%) of all protein class hits (Supplementary Table 10). This was over five-fold enrichment compared to an estimated genome-wide rate (Figure 1d). Notably, 89 immune genes were identified in classical strains, and 84 of these were shared with at least one of the wild-derived strains (Figure 1d). SPRET/EiJ contributed the largest number of strain-specific gene hits (22 genes), whereas WSB/EiJ was the only strain category in which no strain-specific gene hits were identified.

Many paralogous gene families were represented among the hSNP regions (Supplementary Data 3), including genes with functional human orthologs. Several prominent examples include *Apolipoprotein L* alleles; variants of which may confer resistance to *Trypanosoma brucei,* the primary cause of human sleeping sickness^36,37^, IFI16 (Interferon Gamma Inducible Protein 16, member of AIM2-like receptors), a DNA sensor required for death of lymphoid CD4 T cells abortively infected with HIV^38^, NAIP (NLR family apoptosis inhibitory protein) in which functional copy number variation is linked to increased cell death upon *Legionella pneumophila* infection^39^, and secretoglobins (Scgb members) which may be involved in tumour genesis, invasion and suppression, in both human and mouse^40,41^. Large gene families in which little functional information is known were also identified. A cluster of approximately 50 genes, which includes hippocalcin-like 1 *(Hpcal1)* and its homologues, was identified (Chr12:18-25Mbp). *Hpcal1* belongs to the neuronal calcium sensors, expressed primarily in retinal photoreceptors or neurons, and neuroendocrine cells^42^. This region is enriched for hSNPs in all strains excluding C57BL/6J and C57BL/6NJ. Interestingly, within this region, *Cpsf3* (21.29Mbp) is located on an island of high conservation in all strains, and a homozygous knockout of this gene produces subviable lines of C57BL/6NJ mice^43^. Additional examples include another region on chromosome 12 (87-88Mbp) containing approximately 20 eukaryotic translation initiation factor 1A *(eIF1a)* homologs, and on chromosome 14 (41-45Mbp) containing approximately 100 Dlg1-like genes. Genes within all hSNP candidate regions have been identified and annotated (Supplementary Figure 5).

Retrotransposons, RNA mediated mobile genetic elements flanked by long terminal repeats (LTRs), and non-LTR forms such as long interspersed nuclear elements (LINEs) and non-autonomous short interspersed nuclear elements (SINEs) account for approximately 96% of TEs present in the mouse genome^4,44^. We examined retrotransposon content in hSNP dense regions on GRCm38, and compared to an estimated null distribution (1 million simulations), found that the hSNP regions are significantly enriched for both LTRs (empirical p < 1×10^−7^) and LINEs (empirical p < 1×10^−7^) (Supplementary Tables 11,12). Gene retrotransposition has long been implicated in the creation of gene family diversity^45^, novel alleles conferring positively selected adaptations^46^. Once transposed, TEs accumulate mutations over time as the sequence diverges^47,48^. For LTRs, LINEs and SINEs, the mean percent sequence divergence was significantly lower (p < 1×10^−22^) within hSNP regions compared to the rest of the genome (Figure 1e). The largest difference in mean sequence divergence was between LTRs within and outside of hSNP dense regions. Examining only repeat elements with less than 1% divergence (i.e. recent copies), we found these regions are significantly enriched for LTRs (empirical p < 1×10^−7^) and LINEs (empirical p = 0.047). Interestingly, SINEs of all ages were significantly (empirical p = 0.018), and young SINEs marginally (p = 0.049), underrepresented at our candidate loci. SINEs constitute 8.3% of the mouse genome^4^ and may have a role in mutational processes^49^, however the exact role SINEs may play at regions of high polymorphic diversity, and recombination hotspots remains unclear.

### *De novo* assembly of complex gene families

The hSNP scan highlighted multicopy gene families involved in immunity and sensory functions. For the first time, the extensive variation at these loci has been revealed by our *de novo* assemblies, particularly between wild-derived strains. Our data elucidated copy number variation previously unknown in the mouse strains, and uncovered gene expansions, contractions and novel members (<80% sequence identity). For example, 23 distinct clusters of olfactory receptors (ORs) were identified indicating substantial variation among the inbred strains. Olfactory receptors are a large family of genes that include more than 1,200 members annotated in the mouse reference^50^. The human OR repertoire is known to vary between individuals^51^, and presence or absence of particular ORs determine an individual’s ability to perceive certain chemical odours^52^. In mouse, phenotypic differences, particularly diet and behaviour, have been linked to distinct OR repertoires^53,54^, although very little is known about OR diversity between inbred laboratory strains. To this end, we have characterised the CAST/EiJ OR repertoire using our *de novo* assembly and identified 1,249 OR genes (Supplementary Data 5). Relative to the reference strain (C57BL/6J), CAST/EiJ has lost 20 ORs and gained 37 gene family members; 12 novel and 25 supported by published predictions based on mRNA derived from CAST/EiJ whole olfactory mucosa (Figure 2a, Supplementary Table 13)^55^.

**Figure 2:**
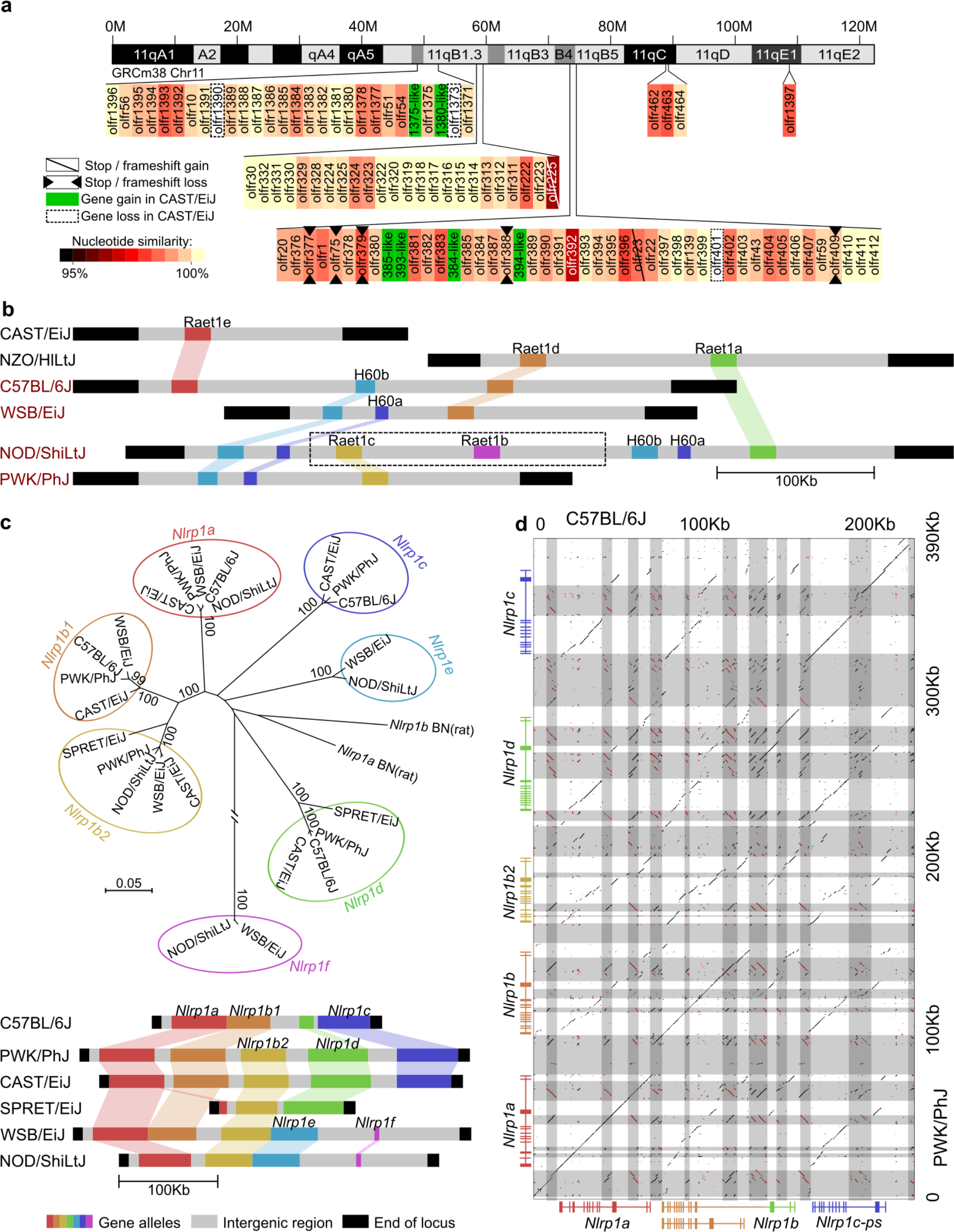
**(a)** Olfactory receptor genes on chromosome 11 of CAST/EiJ. Gene gain/loss and similarity are relative to C57BL/6J. Novel members are named after their most similar homologues. **(b)** Gene order across *Raet1/H60* locus in the collaborative cross parental strains (A/J, NOD/ShiLtJ and 129S1/SvImJ share the same haplotype on this locus, represented by NOD/ShiLtJ). Strain name in black/red indicate *Aspergillus fumigatus* resistant/susceptible. Dashed box indicates unconfirmed gene order. **(c)** Novel proteincoding alleles of the *Nlrp1* gene family in the wild-derived strains and two classical inbred strains. Colours represent the phylogenetic relationships (top, amino acid neighbor joining tree of NBD domain) and the relative gene order across strains (bottom). **(d)** A regional dot plot of the *Nlrp1* locus in PWK/PhJ compared to the C57BL/6J GRCm38 reference (colour-coded same as panel (c)). Grey blocks indicate repeats and transposable elements.

Many gene family loci, particularly immune loci, are characterised by variable copy number among individuals within a population (e.g. MHC^56^). Complete sequencing and annotation of multiple inbred strains in parallel enabled us to accurately predict the underlying structure of many of these gene family loci. We discovered novel gene members at several important immune loci regulating innate and adaptive responses to infection. For example, chromosome 10 (22.1-22.4Mbp) on C57BL/6J contains *Raet1* alleles and minor histocompatibility antigen members *H60*. Raet1/H60 are important ligands for NKG2D, an activating receptor of natural killer cells, and mediate innate immune system detection of infected and stressed cells^57^. These molecules are expressed on the surface of infected^58^ and metastatic cells^59^, and may have a role in allograft autoimmune responses^60^. From the *de novo* assembly, six different *Raet1/H60* haplotypes were identified among the eight Collaborative Cross (CC) founder strains; three of the haplotypes identified are shared among the classical inbred CC founders (A/J, 129S1/SvImJ and NOD/ShiLtJ have the same haplotype), and three different *Raet1/H60* haplotypes were identified in each of the wild-derived inbred strains (CAST/EiJ, PWK/PhJ and WSB/EiJ) (Figure 2b, Supplementary Figure 7). Inspection of the *Raet1/H60* locus using Fiber-FISH fluorescence tags supported our predicted allele structure in all strains (Supplementary Figures 7, 8). The CAST/EiJ haplotype encodes only a single *Raet1* family member *(Raet1e)* and no *H60* alleles, while the classical NOD/ShiLtJ haplotype has four *H60* and three *Raet1* alleles. The *Aspergillus-* resistant locus 4 *(Asprl4),* one of several QTLs that mediate resistance against *Aspergillus fumigatus* infection, overlaps this locus and comprises of a 1MB (∼10% of QTL) interval which, compared to other classical strains, contains a haplotype unique to NZO/HlLtJ (Supplementary Figure 7). Strain specific haplotype associations with *Asprl4* and survival have been reported for CAST/EiJ and NZO/HlLtJ, both of which exhibit resistance to *A. fumigatus* infection^61^. Interestingly, they are also the only strains to have lost *H60* alleles at this locus.

We examined three immunity related polymorphic loci on chromosome 11, *IRG* (GRCm38: 48.85-49.10Mbp), *Nlrp1* (71.05-71.30Mbp) and *Slfn* (82.9-83.3Mbp), because of their polymorphic complexity and importance for mouse survival^62–64^. The *Nlrp1 locus* (NOD-like receptors, pyrin domain containing) encodes inflammasome components that sense endogenous microbial products and metabolic stresses, thereby stimulating innate immune responses^65^. In the house mouse, *Nlrp1* alleles are involved in sensing *Bacillus anthracis* lethal toxin, leading to inflammasome activation and pyroptosis of the macrophage^66,67^. We discovered seven distinct *Nlrp1* family members by comparing six strains (CAST/EiJ, PWK/PhJ, WSB/EiJ, SPRET/EiJ, NOD/ShiLtJ, and C57BL/6J). Each of these six strains exhibit unique haplotype of *Nlrp1* members, highlighting the extensive sequence diversity at this locus across inbred mouse strains (Figure 2c). Each of the three *M. m. domesticus* strains (C67BL/6J, NOD/ShiLtJ and WSB/EiJ) carry different combinations of *Nlrp1* family members; *Nlrp1d-1f* are novel strain-specific alleles discovered that were previously unknown. Diversity between different *Nlrp1* alleles is higher than mouse/rat diversity. For example, C57BL/6J contains *Nlrp1c* which is not present in the other two strains, while *Nlrp1b2* is present in both NOD/ShiLtJ and WSB/EiJ but not C57BL/6J. In PWK/PhJ (*M. m. musculus*), the *Nlrp1* locus is greatly expanded relative to the GRCm38 reference genome, and contains novel *Nlrp1* homologues (Figure 2c), whereas the *Nlrp1* locus is greatly contracted in *M. spretus* (also wild-derived) compared to any other strain. Approximately 90% of intergenic regions in the PWK/PhJ assembly of the locus is composed of transposable elements (Figure 2d).

The wild-derived PWK/PhJ (M. *m. musculus)* and CAST/EiJ (M. *m. castaneus)* strains share highly similar haplotypes, however PWK/PhJ macrophages are resistant to pyroptotic cell death induced by anthrax lethal toxin but CAST/EiJ macrophages are not^68^. It has been suggested that *Nlrp1c* may be the causal family member mediating resistance; *Nlrp1c* can be amplified from PWK/PhJ macrophages but not CAST/EiJ^68^. In the *de novo* assemblies, both mouse strains share the same promoter region for *Nlrp1c;* however, when transcribed, the cDNA of *Nlrp1c_CAST* could not be amplified with previously designed primers^68^ due to SNPs at the primer binding site (5’ …CACT-3’ → 5’ …TACC-3’). The primer binding site in PWK/PhJ is the same as C57BL/6J, however *Nlrp1c* is a predicted pseudogene. We found 18aa mismatches in NBD domains between Nlrp1b_CAST and Nlrp1b_PWK. These divergent profiles suggest that *Nlrp1c* is not the sole mediator of anthrax lethal toxin resistance in mouse, and instead that several other members may also be involved. Newly annotated members *Nlrp1b2* and *Nlrp1d,* appear functionally intact in CAST/EiJ, but were both predicted as pseudogenes in PWK/PhJ due to the presence of stop codons or frameshift mutations. In C57BL/6J, three splicing isoforms of *Nlrp1b* (SV1, SV2, and SV3) were reported^68^. A dot-plot between PWK/PhJ and the C57BL/6J reference illustrates the disruption of co-linearity at the PWK/PhJ *Nlrp1b2* and *Nlrp1d* alleles (Figure 2d). All of the wild-derived strains we sequenced contain a full length *Nlrp1d,* and exhibit a similar disruption of co-linearity at these alleles relative to C57BL/6J (Supplementary Data 6), such that the SV1 isoform in C57BL/6J is derived from truncated ancestral paralogs of *Nlrp1b* and *Nlrp1d,* indicating that *Nlrp1d* was lost in the C57BL/6J lineage. The genome structure of Nlrp1 locus in strains PWK/PhJ, CAST/EiJ, WSB/EiJ and NOD/ShiLtJ were confirmed with Fiber-FISH (Supplementary Figure 9).

The assemblies also revealed extensive diversity at each of the other loci examined; Immunity-related GTPases (IRGs) and Schlafen family (Slfn). IRGs belong to a subfamily of interferon-inducible GTPases present in most vertebrates and throughout the mammalian phylogeny^69^. In mouse, IRG protein family members contribute to the mouse adaptive immune system by conferring resistance against intracellular pathogens such as *Chlamydia trachomatis, Trypanosoma cruzi* and *Toxoplasma gondii*^70^. Our *de novo* assembly is concordant with previously published data for CAST/EiJ^62^, and for the first time has revealed the order, orientation, and structure of three highly divergent haplotypes present in WSB/EiJ, PWK/PhJ, and SPRET/EiJ including novel annotation of rearranged promoters, inserted processed pseudogenes and a high frequency of LINE repeats (Supplementary Data 6).

The Schlafen (Chr11:82.9M-83.3M) family of genes are reportedly involved in immune responses, cell differentiation, proliferation and growth, cancer invasion and chemotherapy resistance, however their exact function remains obscure. In humans, SLFN11 was reported to inhibit HIV protein synthesis by a codon-usage-based mechanism^71^, and in non-human primates positive selection on *Slfn11* has been reported^72^. In mouse, embryonic death may occur between strains carrying incompatible *Slfn* haplotypes^73^. Assembly of Slfn for the three collaborative cross founder strains of wild-derived origin (CAST/EiJ, PWK/PhJ and WSB/EiJ) revealed for the first time extensive variation at this locus. Members of group 4 *Slfn* genes^64^, *Slfn8, Slfn9* and *Slfn10,* show significant sequence diversity among these strains. For example, *Sfln8* is a predicted pseudogene in PWK/PhJ but protein coding in the other strains, although the CAST/EiJ allele contains 78aa mismatches compared to the C57BL/6J reference (Supplementary Figure 10). Both CAST/EiJ and PWK/PhJ contain functional copies of *Sfln10,* which is a predicted pseudogene in C57BL/6J and WSB/EiJ. A novel start codon upstream of *Slfn4,* which causes a 25aa N-terminal extension, was identified in PWK/PhJ and WSB/EiJ. Another member present in the reference, *Slfn14,* is conserved in PWK/PhJ and CAST/EiJ but a pseudogene in WSB/EiJ (Supplementary Figure 10).

### Reference genome updates informed by the strain assemblies

The generation of assemblies from mouse strains closely related to the C57BL/6J reference enabled a new approach to improving the GRCm38 reference assembly. There are currently eleven genes in the GRCm38 reference assembly (C57BL/6J) that are incomplete due to a gap in the sequence. First, these loci were compared to the respective regions in the C57BL/6NJ assembly and used to identify contigs from public assemblies of the reference strain, previously omitted due to insufficient overlap. Second, C57BL/6J reads aligned to the regions of interest in the C57BL/6NJ assembly were extracted for targeted assembly, leading to the generation of contigs covering sequence currently missing from the reference. Both approaches resulted in the completion of ten new gene structures (e.g. Supplementary Figure 11 and Supplementary Data 7), and the near-complete inclusion of the Sts gene that was previously completely missing from the assembly.

Improvements to the reference genome, coupled with pan-strain gene predictions, were used to provide updates to the existing reference genome annotation, maintained by the GENCODE consortium^74^. We examined the strain specific RNA-Seq (Comparative Augustus) gene predictions containing 75% novel introns compared to the existing reference annotation (Table 1) (GENCODE M8, chromosomes 1-12). Of the 785 predictions investigated, 62 led to the annotation of new loci including 19 protein coding genes and 6 pseudogenes (Supplementary Table 14, Supplementary Data 8). In most cases where a new locus was predicted on the reference genome, we identified pre-existing, but often incomplete, annotation. The predictions highlighted unannotated exons and splice sites that could be confirmed with orthogonal supporting data such as intron support from RNAseq experiments. For example, the *Nmur1* gene was extended at its 5’ end and made complete on the basis of evidence supporting a prediction which spliced to an upstream exon containing the previously missing start codon. The *Mroh3* gene, which was originally annotated as an unprocessed pseudogene, was updated to a protein coding gene due to the identification of novel intron that permits extension of the CDS to full-length. The previously annotated pseudogene model has been retained as a nonsense-mediated decay (NMD) transcript of the protein coding locus. At the novel bicistronic locus, *Chml_Opn3,* the original annotation was single exon gene, *Chml*, that was extended and found to share its first exon with the *Opn3* gene.

**Table 1.**
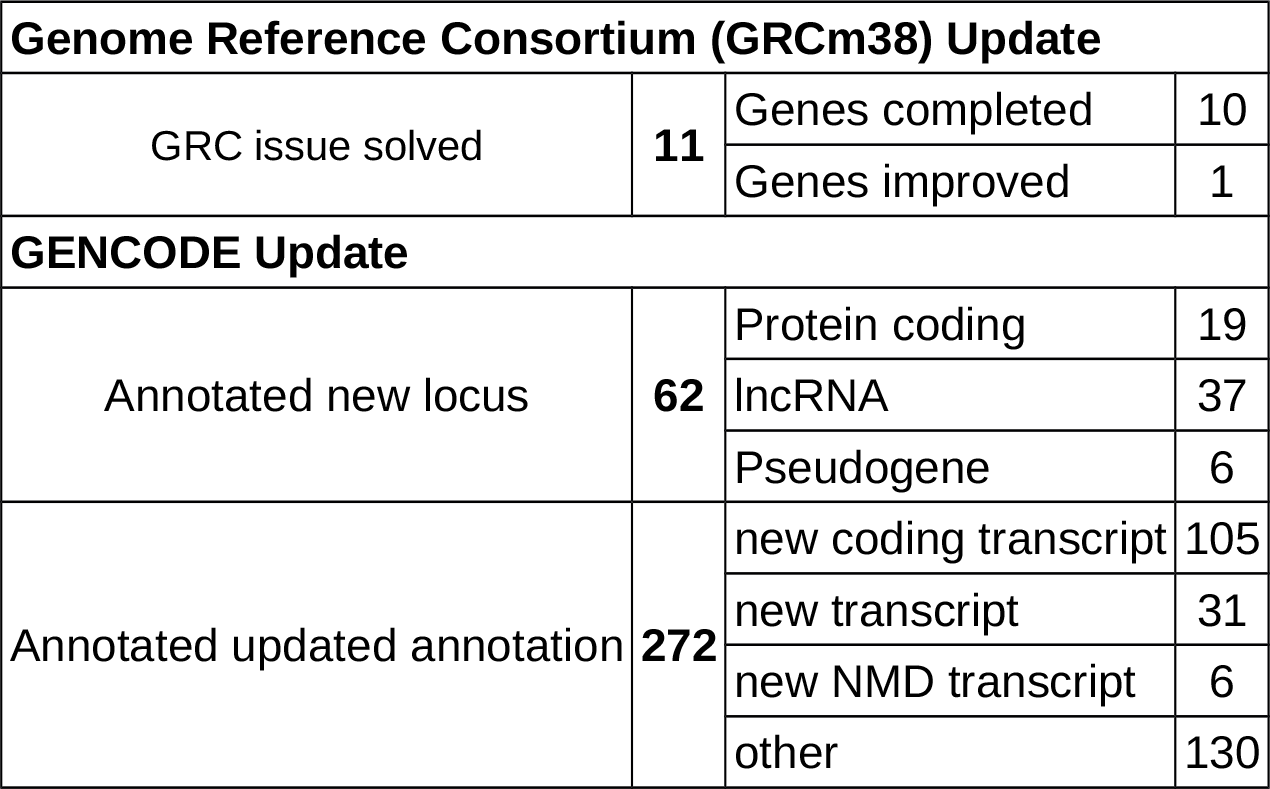
Genome Reference Consortium (GRCm38) and GENCODE annotation updates informed by the strain assemblies. Updates indicate known GRC issues solved based on C57BL/6NJ *de novo* assembly. GENCODE update is based on comparative Augustus predictions with 75% novel introns and includes annotation and predictions which occur on chromosomes 1-12.

We discovered a novel 188-exon gene on chr11 that significantly extends the existing gene *Efcab3* spanning between *Itgb3* and *Mettl2.* This *Efcab3-like* gene was manually curated, validated according to HAVANA guidelines^75^, and identified in GENCODE releases M11 onwards as Gm11639. *Efcab3/Efcab13* are calcium-binding proteins and the new gene primarily consists of repeated EF-hand protein domains (Supplementary Figure 12). Analysis of synteny and genome structure revealed that the *Efcab3* locus is largely conserved across other mammals including most primates. Comparative gene prediction identified the full length version in orangutan, rhesus macaque, bushbaby and squirrel monkey. However, the locus contains a breakpoint at the common ancestor of chimpanzee, gorilla and human *(Homininae)* due to a ∼15Mbp intra-chromosomal rearrangement that also deleted many of the internal EF-hand domain repeats (Figure 3b, Supplementary Figure 13). Analysis of GTEx expression data^76^ in human revealed that the *EFCAB13* locus is expressed across many tissue types, with the highest expression measured in testis and thyroid. In contrast, the *EFCAB3* locus only has low level measurable expression in testis. This is consistent with the promoter of the full length gene being present upstream of the *EFCAB13* version, which is supported by H3K4Me3 analysis (Supplementary Figure 14). In mice, *Efcab3* is specifically expressed during development throughout many tissues with high expression in the upper layers of the cortical plate (source, http://www.genepaint.org), and is located in the immediate vicinity of the genomic 17q21.31 syntenic region linked to brain structural changes both in mice and humans^77^. As such, we used a recently developed and highly robust approach for the assessment of 40 brain parameters across 22 distinct brain structures (Supplementary Table 15), and analysed neuroanatomical defects in *Efcab3-like*^−/−^ mice (see methods). This consisted of a systematic quantification of the same sagittal brain region at Lateral +0.72 mm, down to cell level resolution and blind to the genotype (Supplementary Table 16). To minimize environmental and genetic variation, mice were analysed according to their gender, aged exactly to 16 week old before brain necropsy, and bred on the same genetic background (C57BL/6NJ). Overall size anomalies were identified in the *Efcab3-like*^−/−^ mice with the majority of assessed parameters increased in size when compared to matched wt (Figure 3c). Interestingly, the lateral ventricle was one the most severely affected brain structure exhibiting an enlargement of 65% (P=0.007). The pontine nuclei were also increased in size by 42% (P=0.001) and the cerebellum by 27% (P=0.02), these are two regions involved in motor activity (Figure 3d). The thalamus was also larger by 19% (P=0.007). As a result, the total brain area parameter was enlarged by 7% (P=0.006). Taken together, these results suggest a mechanism of *Efcab3-like* to regulate brain development and brain size regulation from the forebrain to the hindbrain.

**Figure 3:**
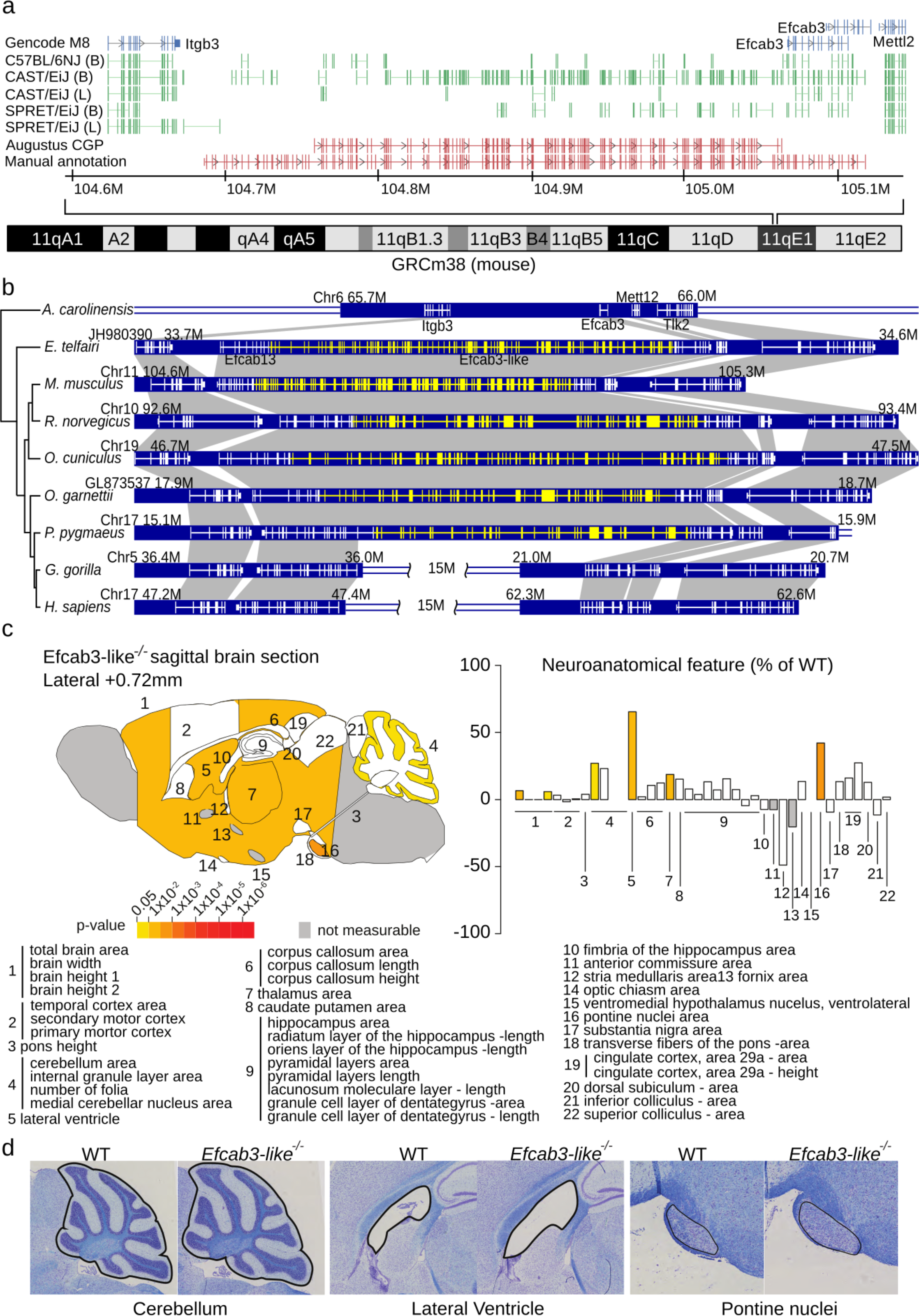
**(a)** Comparative Augustus identified a previously unannotated 188 exon gene *(Efcab3-like,* red tracks) present in all strains. RNA-Seq splice sites from two tissues (B=Brain, L=Liver, green tracks) and five strains are displayed. Manual annotation extended this novel gene to 188 exons (lower red track). **(b)** Evolutionary history of *Efcab3-like* in vertebrates and genome structure of *Efcab3-like* and surrounding genes. The mRNA structure of each gene is shown with white lines on the blue blocks and novel coding sequence discovered in this study is shown in yellow. Notably, both *Efcab13* and *Efcab3* are parts of novel gene *Efcab3-like.* A recombination event happened in the common ancestor of sub-family *Homininae,* which disrupted Efcab3-like in gorilla and chimpanzee (not shown) human. **(c)** Schematic representation of 22 unique brain regions plotted in sagittal plane at Lateral +0.72 mm of the Mouse Brain Atlas^91^ for *Efcab3-like∼^l^∼* male mice (16 weeks of age) according to p-values (left). Corresponding brain regions are labelled with a number that is described below the panel (Supplementary Table 15). White colouring indicates a p-value > 0.05 and grey indicates that the brain region could not be confidently tested due to missing data. Raw neuroanatomical data are available in Supplementary Table 16. Histograms showing the neuroanatomical features as percentage increase or decrease of the assessed brain regions in *Efcab3-like∼^l^∼* mice as compared to the matched controls (100%) at Lateral +0.72 mm (right). **(d)** Representative sagittal brain images, double-stained with luxol fast blue and cresyl violet acetate, of matched controls (left) and *Efcab3-like∼^l^∼* (right), showing a larger cerebellum, enlarged lateral ventricle and increased size of the pontine nuclei.

## Discussion

The completion of the mouse reference genome, based on the classical inbred strain C57BL/6J, was a transformative resource for human and mouse genetics. There are many other mouse strains that offer a rich source of genetic and phenotypic diversity in widespread use^8^. We generated the first chromosome scale genome assemblies for 12 classical and 4 wild-derived inbred strains, thus revealing at unprecedented resolution the striking strain-specific allelic diversity that encompasses 0.5-2.8% (14.4-75.5 Mbp, excluding C57BL/6NJ) of the mouse genome. Accessing shared and distinct genetic information across the *Mus* lineage in parallel during assembly and gene prediction leads to the placement of novel alleles, the accurate annotation of many strain-specific gene family haplotypes, and the detection of genes lowly expressed but partially supported in all strains (Figure 3a). Our *de novo* assembly revealed novel genome diversity between mouse strains especially in the wild-derived mice. This is particularly important for experimental studies involving inbred laboratory strains, whose response to an experimental condition (such as an infection or diet) may be contingent on presence or absence of individual genes. Many regions exhibit gene family and sequence diversity in the strains, including founders of key recombinant inbred mouse panels of both classical and wild-derived origin, and contain novel members at previously reported response loci. Mouse recombinant inbred and outbred panels (e.g. CC, DO, HS) are commonly used to investigate mammalian physiology, gene function during infection, inheritance and disease networks^11,78^. Although progeny are derived with knowledge of parental origins, previously the underlying genome structure of the founders at many loci has remained elusive. Larger genomic events, including duplications, and novel members not present in the reference can be difficult to quantify through variant identification alone. This is particularly prevalent at loci that exhibit extreme variation from the reference, where different gene combinations, even among classical strains, are a common feature of many gene families and complete subsets of alleles are not represented (e.g. Figure 2a). Alignment to the true contributing haplotype, or a more closely related reference genome, may facilitate improved annotation of disease variants, and even localise responses to individual gene family members unique to individual strains.

Genetic diversity at gene loci, particularly those related to defence and immunity, is often the result of selection that if retained, can lead to the rise of divergent alleles in a population^79^. This can be the result of host-pathogen interactions but examples of other diverse expansions have been observed in evolutionary lineages^80^. Many protein coding genes are known to have undergone recent lineage specific expansion in mouse^5^ (e.g. *Abp*^81^, *Rhox*^82^,and *Mups*^83^), and appear to be retained by balancing selection (e.g. *Oas1b*^84^, *IRG*^62^). Perturbing gene copy number can give rise to clusters of genes usually with similar sequence and function, often composed of unique combinations from the entire family repertoire. We used the presence of dense clusters of heterozygous SNPs on the C57BL/6J reference genome as a marker for extreme polymorphism, and examine the *de novo* assembly to explore the underlying architecture of the lineage specific changes. These account for between 1.5-5.5% of protein coding genes (Figure 1c) and are overrepresented for immunity, sensory, sexual reproduction, and behavioral phenotypes (Figure 1d). Genes related to immunological processes, particularly gene families involved in mediating innate immune responses (e.g. *Raet1, Nlrp1),* exhibit great diversity among the strains reflecting strain-specific disease associations, responses and susceptibility. Interestingly, regions of strain haplotype diversity appear enriched for recent LINEs and LTR repeat elements (Figure 1e). Retrotransposons have long been implicated in evolution of gene function in mammalian genomes, and can foster adaptive responses to selective pressure by facilitating recombination within a population leading to increased allelic diversity^85^, and are a key element of population fitness and pathogen resistance^86,87^. Indeed, we observed several innate immunity gene families in mice with high density of retrotransposons, which is the likely mechanism for diversification at that gene locus (e.g. *Nlrp1,* Figure 2d).

Having access to multiple chromosome scale genome sequences from within a species is the foundation for understanding the genetic mechanisms of phenotypic differences and traits. The challenge of generating multiple closely related mammalian genomes and annotation required new approaches to whole-genome alignment^88^, comparative creation of whole-chromosome scaffolds^89^, and comparative approaches to simultaneous genome annotation within a clade^30,31^. *Mus* is the first mammalian lineage to have multiple chromosome scale genomes. Simultaneous access to many rodent species assemblies in parallel with individual level gene predictions, expression and long read data facilitated the accurate prediction of many strain specific haplotypes and gene isoforms. This approach identified previously unannotated genes, including *Efcab3-like*, one of the largest known mouse genes (5874aa) which also appears conserved in mammals. Interestingly, the previously unannotated Efcab3-like gene is very close to the 17q21.31 syntenic region associated in humans to the Koolen-de Vries microdeletion syndrome (KdVS). Both mouse deletion models of this syntenic interval^77^, containing four genes *(Crhr1, Spplc2, Mapt* and *Kansl1*; Figure 3a) and *Efcab3-like* knockout showed analogous brain phenotypes, suggesting common cis-acting regulatory mechanisms as shown previously in the context of the 16p11.2 microdeletion syndrome^90^. *Efacb3-like* is conserved in orangutan but reversed in gorilla, and appears to have split into two separate protein coding genes, *EFCAB3* and *EFCAB13*, in the *Homininae* lineage. Many novel genes and strain specific transcripts were identified across all of the strains, highlighting unexplored sequence variation across the *Mus* lineage. The addition of these genomes, in particular C57BL/6NJ, enabled the resolution of GRCm38 reference assembly issues, and the improvement of several reference gene annotations. Strain specific gene annotations are critical to understand inheritance patterns, and examine the effect of variation, haplotype structure and gene combinations associated with disease. In particular, the wild-derived strains represent a rich resource of novel target sites, resistance alleles, genes and isoforms not present in the reference strain, or indeed many classical strains. For the first time the underlying sequence at these loci is represented in strain-specific assemblies and gene predictions from across the inbred mouse lineage, which should facilitate increased dissection of complex traits.

## Methods

See separate online materials and methods document.

## Data availability

The genome sequencing reads are available from the European Nucleotide Archive and the assemblies are part of NCBI BioProject PRJNA310854 (Supplementary Table 17). The genome assemblies and annotation are available via the Ensembl genome browser, and the UCSC genome browser. Sequence accessions for the three immune related loci on Chr11 are available from the European Nucleotide Archive (Supplementary Table 18).

## Acknowledgements

This work was supported by the Medical Research Council [MR/L007428/1], BBSRC [BB/M000281/1] and the Wellcome Trust. DJA is supported by Cancer Research-UK and the Wellcome Trust. We thank members of the Sanger Institute Mouse Pipelines teams (Mouse Informatics, Molecular Technologies, Genome Engineering Technologies, Mouse Production Team, Mouse Phenotyping) and the Research Support Facility for the provision and management of the mice. We thank Valerie Vancollie for assistance with phenotyping data.

